# Structural variability and concerted motions of the T cell receptor – CD3 complex

**DOI:** 10.1101/2021.02.03.429549

**Authors:** Prithvi R. Pandey, Bartosz Różycki, Reinhard Lipowsky, Thomas R. Weikl

## Abstract

We investigate the structural and orientational variability of the membrane-embedded T cell receptor (TCR) – CD3 complex in extensive atomistic molecular dynamics simulations based on the recent cryo-EM structure determined by ***Dong et al.*** (***2019***). We find that the TCR extracellular (EC) domain is highly variable in its orientation by attaining tilt angles relative to the membrane normal that range from 15° to 55°. The tilt angle of the TCR EC domain is both coupled to a rotation of the domain and to characteristic changes throughout the TCR – CD3 complex, in particular in the EC interactions of the Cβ FG loop of the TCR, as well as in the orientation of transmembrane helices. The concerted motions of the membrane-embedded TCR – CD3 complex revealed in our simulations provide atomistic insights for force-based models of TCR activation, which involve such structural changes in response to tilt-inducing forces on antigen-bound TCRs.

## Introduction

T cells recognize peptide antigens presented by major histocompatibility complexes (MHC) on apposing cell surfaces as a central step in the initiation of adaptive immune responses (***Rossjohn et al., 2015***; ***Smith-Garvin et al., 2009***; ***Dustin, 2014***; ***Pettmann et al., 2018***; ***Belardi et al., 2020***). The antigen recognition is performed by the T-cell receptor (TCR) complex, a complex of four dimeric transmembrane proteins. In this complex, the heterodimeric TCRαβ contains the binding site for recognizing peptide antigens, and the associated CD3єδ and CD3єγ heterodimers and the CD3*ζ ζ* homodimer contain the intracellular signaling motifs that transmit antigen binding to T cell activation (***Wucherpfennig et al., 2010***). While this stoichiometry of the complex has been known for nearly two decades (***Call et al., 2002***), the structure of the TCR – CD3 complex remained a puzzling problem (***Fernandes et al., 2012***; ***Birnbaum et al., 2014***; ***Natarajan et al., 2016***) that has only been recently solved by ***Dong et al. (2019***) with cryogenic electron microscopy (cryo-EM). To determine the structure, Dong et al. expressed all proteins of the complex in cultured cells, replaced the cell membrane around the assembled TCR – CD3 complex by the detergent digitinon, and stabilized the interactions between the extracellular (EC) domains of TCRαβ, CD3єδ and CD3єγ by chemical crosslinking. In the cryo-EM structure, the EC domains of CD3єδ and CD3єγ are both in contact with TCRαβ and with each other (see Figure 1), which explains the cooperative binding of CD3єδ and CD3єγ to TCRαβ observed in chain assembly (***Call et al., 2002***), mutational (***Kuhns and Davis, 2007***, ***2012***), and NMR experiments (***He et al., 2015***). As indicated by mutational experiments (***Kuhns and Davis, 2007***, ***2012***), the DE loop of the membrane-proximal constant domain Cα of TCRα is in contact with the CD3єδ EC domain, and the CC’ loop of the constant domain Cβ of TCRβ is in contact with both the CD3єγ and CD3єδ EC domains in the assembled TCR – CD3 complex (see Figure 1). Outstanding questions concern the orientational variability of the TCRαβ EC domain relative to the membrane, in which the TCR – CD3 complex complex is embedded in its native environment, and the structural variability of the overall TCR – CD3 complex, which is constrained by the chemical crosslinking of the protein chains in the approach of Dong et al.(***Dong et al., 2019***; ***Reinherz, 2019***). The Cβ FG loop, for example, has been suggested to play a key role in T cell activation (***Kim et al., 2010***; ***Touma et al., 2006***), but exhibits only rather limited contacts with CD3єγ in the cryo-EM structure (see Figure 1).

**Figure 1.**
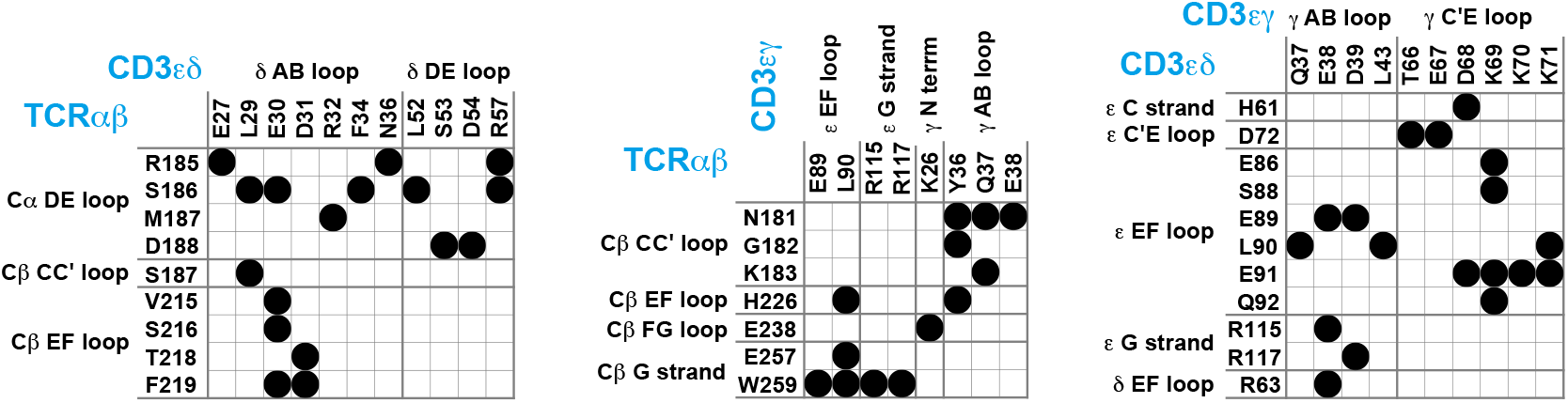
Maps of residue-residue contacts (black disks) between the EC domains of the protein dimers TCRαβ, CD3єδ, and CD3єγ in the cryo-EM structure of the T cell receptor – CD3 complex (Dong et al., 2019). Here, two residues are taken to be in contact if the minimum distance between non-hydrogen atoms of the residues is smaller than 0.45 nm. The loops and strands of the membrane-proximal constant domains Cα and Cβ of the proteins TCRα and TCRβ and of the EC domains of CD3є, CD3γ, and CD3δ are labeled according to the standard convention for immunoglobulin-like domains (***Garcia et al., 1996***; ***Wang et al., 1998***; ***Sun et al., 2004***).

In this article, we investigate the structural and orientational variability of the membrane-embedded TCR – CD3 complex in extensive, atomistic molecular dynamics (MD) simulations with a cumulative simulation length of 120 μs. Compared to the cryo-EM structure, significantly more residues of TCRαβ are involved in EC domain contacts along our simulation trajectories, in particular in the Cβ FG loop, and notably also in the variable domain Vα of the TCRα chain. We 1nd that the TCRαβ EC domain is rather variable in its orientation, with tilt angles relative to the membrane normal that range from 15° to 55°. The tilt of the TCRαβ EC domain is both coupled to a rotation of the domain and to characteristic changes in the overall structure of the TCR – CD3 complex. These structural changes include a clear decrease of contacts in the Cβ FG loop and an increase of contacts in the Vα domain with increasing tilt angle of the TCRαβ EC domain, as well as changes in the orientation of the transmembrane (TM) helices of the TCRα and CD3γ chain. The concerted motions of the membrane-embedded TCR – CD3 complex revealed in our simulations provide atomistic insights for force-based models of TCR signalling, which involve structural changes, in particular in the Cβ FG loop, that are induced by transversal, tilt-inducing forces on bound TCRs (***Brazin et al., 2015***; ***Feng et al., 2018***).

## Results

Our computational analysis of the structural and orientational variability of the membrane-embedded TCR – CD3 complex is based on 120 atomistic, explicit-water MD simulation trajectories with a length of 1 μs and, thus, on simulation data with a total length of 120 μs. We have conducted these simulations with the Amber99SB-ILDN protein force field (***Lindorff-Larsen et al., 2010***) and the Amber Lipid14 membrane force field (***Dickson et al., 2014***) at a simulation temperature of 30°C on graphics processing units (GPUs). The simulation trajectories start from initial system conformations in which the cryo-EM structure of the TCR – CD3 protein complex is embedded in a membrane composed of 456 POPC lipids and 114 cholesterol molecules. We find that the orientational and conformational ensembles sampled by the 120 trajectories equilibrate within the first 0.5 μs of the simulation trajectories (see Methods) and, therefore, focus on the second 0.5 μs of the MD simulation trajectories in our analysis.

Compared to the cryo-EM structure, a much larger set of residues is involved in contacts between the protein dimers of the TCR – CD3 complex in our MD simulations. Figure 2 illustrates the time-averaged contacts between residues of the TCRαβ, CD3єγ, and CD3єδ EC domains along the equilibrated second halves of the MD simulation trajectories. The fourth protein dimer in the complex, CD3*ζ ζ*, has no EC domain. The contact maps of Figure 2 include all residue-residue contacts that occur in the simulations with a probability larger than 0.5%. The probabilities of the contacts are calculated from 6000 simulation structures extracted at intervals of 10 ns from the second 0.5 μs of the 120 simulations, and indicated in grayscale in Figure 2. The contacts are grouped in clusters (blue numbers) that correspond to interactions between loops and strands of the EC domains, which are labeled according to the standard convention for immunoglobulin(Ig)-like domains (***Garcia et al., 1996***; ***Wang et al., 1998***; ***Sun et al., 2004***). The EC domains of the proteins TCRα and TCRβ consist of the membrane-proximal constant domains Cα and Cβ and the variable domains Vα and Vβ, which are all Ig-like domains, as are the EC domains of the proteins CD3є, CD3γ, and CD3δ. In our MD simulations, significantly more loops and strands, and more residues of the protein dimers TCRαβ, CD3єγ, and CD3єδ participate in EC domain interactions, compared to the cryo-EM structure. The Cβ FG loop, for example, exhibits only a single contact with an N-terminal residue of CD3γ in the cryo-EM structure (see Figure 1). In our MD simulations, in contrast, the Cβ FG loop is involved in a large number of contacts with the N-terminus of CD3γ and with several loops and strand in the є chain of the CD3єγ EC domain. Besides the Cα DE loop, Cβ CC’ loop, Cβ EF loop, Cβ FG loop, and Cβ G strand with contacts in the cryo-EM structure, the MD contacts maps of Figure 2 include also the Cα AB loop and the Cβ A and B strand in the constant domains of TCRαβ and, remarkably, the three loops A’B, C”D, and EF in the variable region Vα of TCRα. Residue-residue contacts between Vα and the δ chain of the CD3єδ have probabilities smaller than 3%, but occur in 75 of the 120 trajectories and are, thus, a robust feature of our simulations. These contacts are grouped in four small, correlated contact clusters (see cluster-cluster correlation coeZcients in Figure 2–figure supplement 1).

**Figure 2.**
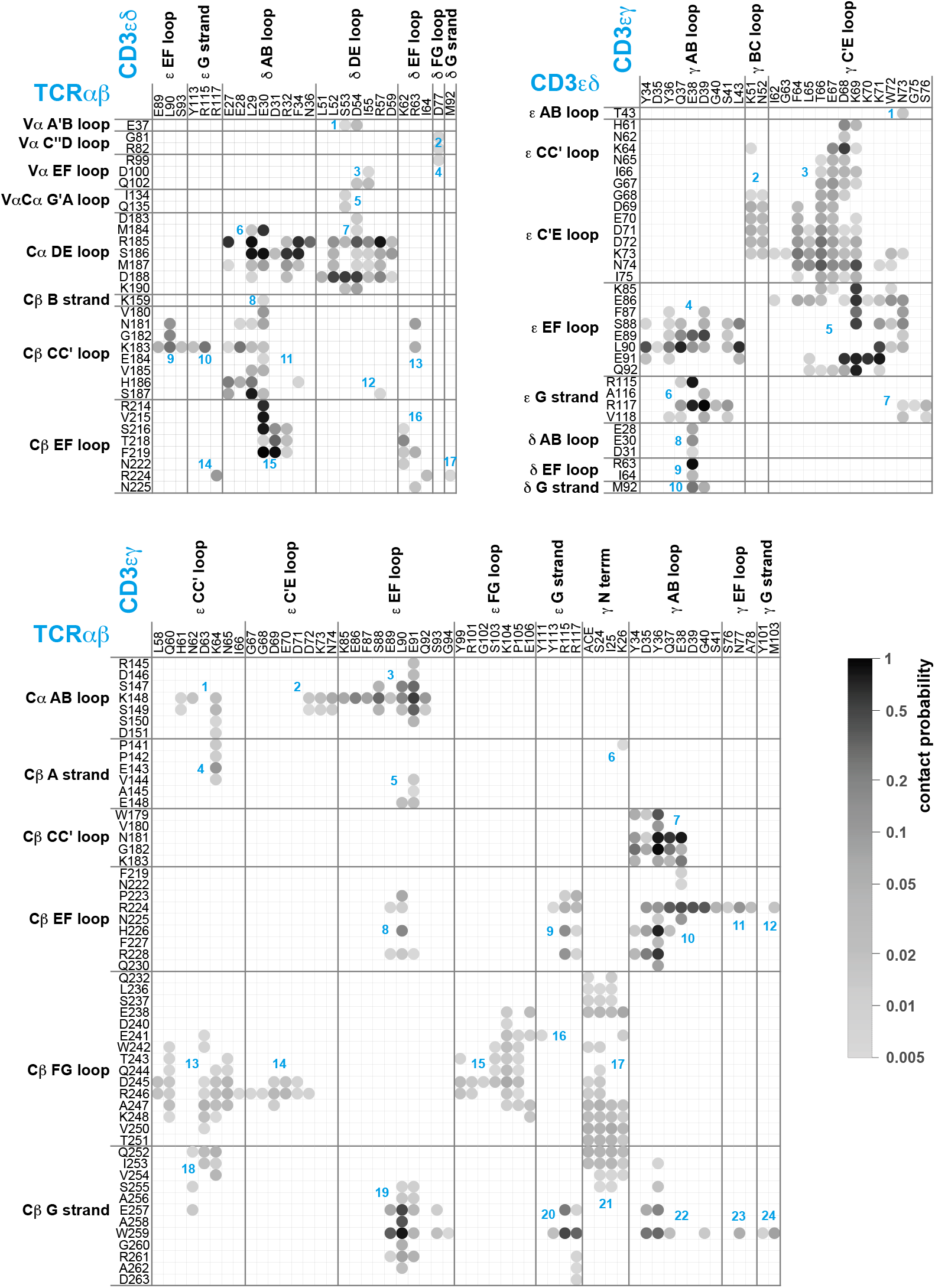
Averaged maps of contacts between the EC domains of TCRαβ, CD3єδ, and CD3єγ in the MD simulation trajectories. The shading of the contact disks indicates the contact probability, i.e. the fraction of simulation structures in which the contact is present. The contact analysis is based on 120 × 50 = 6000 structures extracted at intervals of 10 ns from the second halves of the 120 microsecond-long trajectories, which reflect an equilibrated ensemble of simulation conformations (see Methods) and are available at the Edmond Open Research Data Repository (***Pandey and Weikl, 2021***). For clarity, only contacts with a contact probability larger than 0.5% are represented. As in Figure 1, two residues are taken to be in contact in a simulation structure if the minimum distance between non-hydrogen atoms of the residues is smaller than 0.45 nm. The contacts occur in clusters with numbers labelled in blue.

In our MD simulations, the TCRαβ EC domain is rather variable in its orientation relative to the membrane. The orientation can be quantified by two angles, a tilt angle and a rotation angle. To determine these angles, we choose two axes A and B in the TCRαβ EC domain: Axis A connects the centres of mass of Cαβ and Vαβ, where Cαβ is the dimer of the constant domains Cα and Cβ, and Vαβ is the dimer of the variable domains Vα and Vβ. Axis B connects the centres of mass of the variable domains Vα and Vβ. The tilt angle of the TCRαβ EC domain then is the angle between axis A and the membrane normal, and the rotation angle is the angle between axis B and the normal of the plane spanned by axis A and the membrane normal. The rotation angle describes the rotation of the TCRαβ EC domain around axis A (see Figure 3(a) and (b)). In our MD simulations, the tilt angle of TCRαβ EC domain roughly varies between 15°and 55°, while the rotation angle varies between 0° and 55°. The two-dimensional probability distribution of the angles in Figure 3(c) indicates that the rotation of the TCRαβ EC domain is coupled to its tilt (see Figure 3(c)): For tilt angles between 15° and 35°, the rotation angle predominantly adopts values between about 5° and 25°. For a tilt angle of 40°, the most probable value of the rotation angle is about 32°, and further increases to 40° for a tilt angle of 50°. The coupling between the tilt and rotation of the TCRαβ EC domain is also illustrated in Figure 3(a) and (b). In the structure of the membrane-embedded TCR – CD3 complex of Figure 3(a), the TCRαβ EC domain has a tilt angle of 32.8° and a rotation angle of 12.8°. In the structure of Figure 3(b) with a larger tilt angle of 50.8°, the rotation angle of the TCRαβ EC domain is 42.9°.

**Figure 3.**
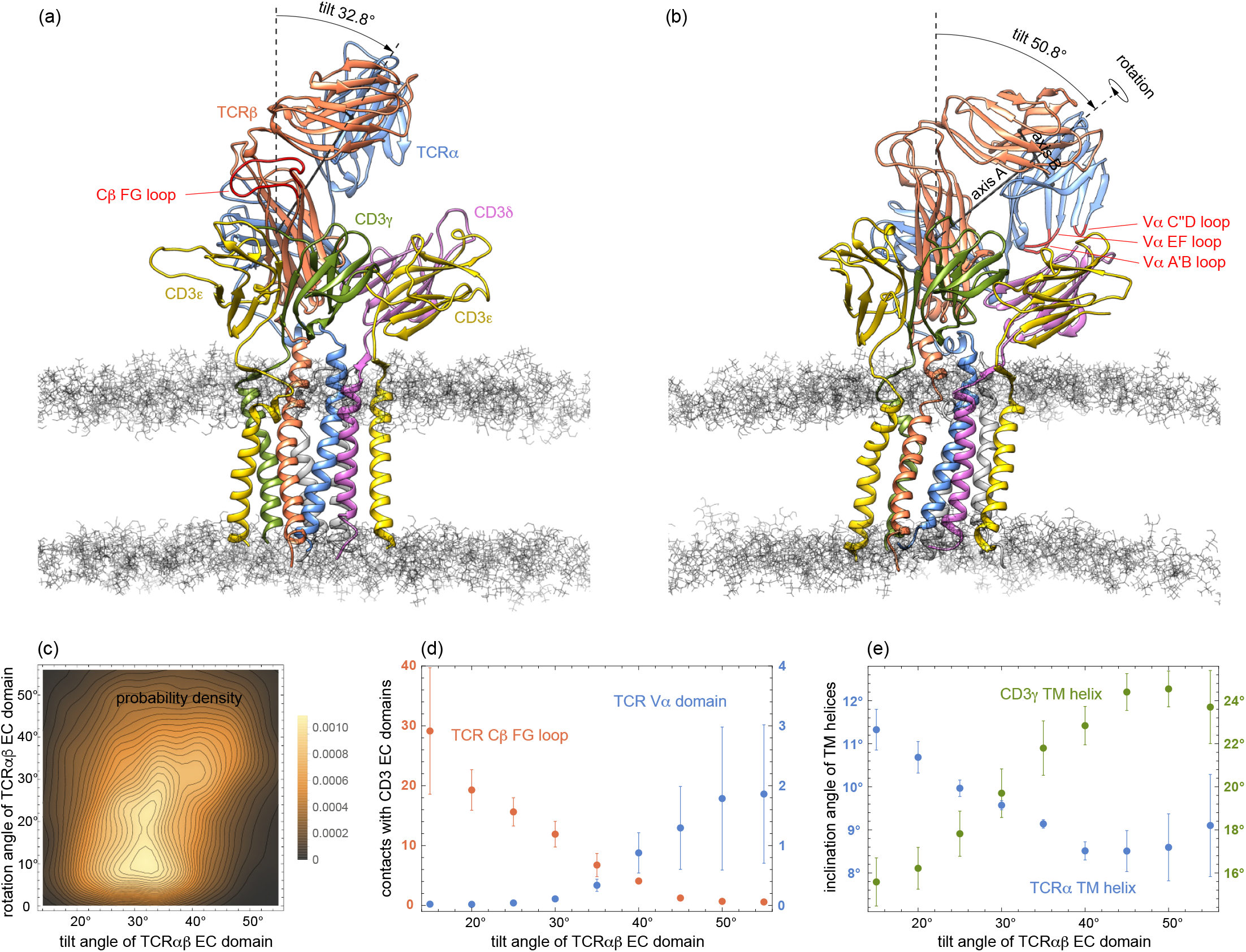
(a) and (b) MD conformations of the TCR – CD3 complex with different tilt angles of the TCRαβ ECdomain relative to the membrane normal. The rotation angles of the TCRαβ ECdomain are 12.8° and 42.9° in the conformations (a) and (b), respectively. (c) Two-dimensional probability density function for the tilt angle and rotation angle of the 6000 equilibrated MD conformations from the 120 trajectories. (d) Numbers of residue-residue contacts with CD3 EC domains for the Cβ FG loop and the Vα domain versus tilt angle. (e) Inclination angle of the TM helices in the TCRα and CD3γ chain relative to the membrane normal as a function of the tilt angle of the TCRαβ ECdomain.

The tilt angle of the TCRαβ EC domain is associated with characteristic changes in the overall structure of the TCR – CD3 complex, in particular with changes in the number of residue-residue contacts of the Vα domain and of the Cβ FG loop (see Figure 3(d)) and in the orientations of the transmembrane (TM) helices of the TCRα and CD3γ chains (see Figure 3(e)). Residue-residue contacts of the A’B, C”D, and EF loops of the variable domain Vα with the protein CD3δ only occur for tilt angles of the TCRαβ EC domain larger than about 30° (see Figure 3(d)). The average number of these residues-residue contacts increases to values around 2 for tilt angles of 50° and larger. For the Cβ FG loop, in contrast, the average number of residues-residue contacts decreases from a value around 30 at the tilt angle 15° to values close 0 for tilt angles of 50° and larger. A decrease in the average number of contacts with increasing tilt angle can also be observed for the Cα AB loop and the Cβ A strand, CC’ loop, and G strand (see Figure 3–figure supplement 1). Only for the Cα DE loop and Cβ EF loop, the average number of contacts is rather independent of the tilt angle. The tilt of the TCRαβ EC domain also affects the orientation of TM helices. The average inclination of the TCRα TM helix relative to the membrane normal decreases from about 11.5° to values around 8.5° with increasing tilt angle of the TCRαβ EC domain, while the average inclination of the CD3γ TM helix increases from about 15.5° to values around 24° (see Figure 3(e)). An increase from about 19° to values around 22° with increasing tilt angle of the TCRαβ EC domain also occurs for the average inclination angle of the TM helix of the є chain of CD3єγ (see Figure 3–figure supplement 2). The average orientation of the other 1ve TM helices relative to the membrane normal exhibits only small variations with the tilt angle of the TCRαβ EC domain.

## Discussion

The coupling of the tilt angle of the TCRαβ EC domain to overall conformational changes in the TCR – CD3 complex, which we observe in our MD simulations, is of relevance for TCR signaling mechanisms that are based on transversal, tilt-inducing forces. Transversal forces acting on the TCRαβ EC domain after binding to MHC-peptide-antigen complexes arise during the scanning of antigen-presenting cells by T cells (***Göhring et al., 2020***; ***Cai et al., 2017***; ***Huse, 2017***; ***Rushdi et al., 2020***). While experiments show that the TCR – CD3 complex responds to mechanical force (***Kim et al., 2009***; ***Feng et al., 2017***) and that the Cβ FG loop plays a key role in this response (***Das et al., 2015***), an outstanding question is how this force alters the conformation of the TCR – CD3 complex (***Courtney et al., 2018***). Our MD simulations show that an increased tilt of the TCRαβ EC domain leads to a marked decrease in the contacts between the Cβ FG loop and the CD3єγ EC domain (see Figure 3(d)), and also to changes in the inclination of TM helices relative to the membrane normal (see Figure 3(e)). Such structural changes in the TM domain of the TCR – CD3 have been suggested to be involved in the transmission of forces from the EC domain to the signaling motifs on the intracellular segments of the CD3 chains (***Brazin et al., 2015***, ***2018***), which are not resolved in the cryo-EM structure. In alternative mechanisms, a key step in T cell activation is the size-based segregation of the inhibitory tyrosine phosphatase CD45 from close-contact zones in which TCR – CD3 complexes can bind to MHC-peptide-antigens on opposing cell surfaces (***Davis and van der Merwe, 2006***; ***Choudhuri and van der Merwe, 2007***; ***Chang et al., 2016***).

The concerted conformational changes in the TCR – CD3 complex are also reflected in the correlations of contact clusters in the EC domain interactions. The largest positive correlations of clusters in the TCRαβ/CD3єδ contact map of Figure 2 occur between the contact clusters 1 to 4 (see Figure 2–figure supplement 1, top left), which correspond to interactions of the variable domain Vα of TCRα and the δ chain of CD3єδ. The large positive correlations of these four contact clusters can be understood from the coupling to the tilt of the TCRαβ EC domain, because the contacts of the Vα domain reported by the four contact clusters are only possible at high tilt angles of the TCRαβ EC domain (see Figure 3(d)). Relatively large positive correlations occur also between the clusters 13 to 21 of the TCRαβ/CD3єγ contact map (see Figure 2–figure supplement 1, bottom left). These clusters reflect interactions of the Cβ FG loop and G strand with the є chain and the N-terminus of the γ chain of CD3єγ, which decrease with increasing tilt angle of of the TCRαβ EC domain (see Figures 3(d) and 3–figure supplement 1). Besides these positive correlations that result from the concerted conformational changes coupled to the tilt angle of the TCRαβ EC domain, the overall weak correlations between the majority of the other contact clusters in Figure 2 indicate independent motions of the loops and strands involved in these EC domain contacts. In addition, relatively strongly negative correlations of several pairs of contact clusters point to alternative EC domain contacts. For example, the overall rather negative correlations between the clusters 13 to 21 and the cluster 22 of the TCRαβ/CD3єγ contact map show that the interaction of the Cβ G strand and the γ AB loop reported by cluster 22 is not compatible with the interactions of the Cβ FG loop and G strand to the є chain and the N-terminus of the γ chain of CD3єγ, which are reflected by the clusters 13 to 21.

Overall, our simulations reveal that the orientation of TCR EC domain relative to the membrane is coupled to structural changes throughout the TCR – CD3 complex. Besides this concerted structural motion, the overall weak correlations of the majority of EC domain interactions indicate additional independent motions of the loops and strands that are involved in these interactions.

## Methods

### System setup

To embed the cryoEM structure of the human TCR-CD3 complex (PDB ID 6jxr) into a lipid membrane, we have first aligned the protein complex along the *z*-axis of the simulation box. In this alignment with the program Visual Molecular Dynamics (VMD) (***Humphrey et al., 1996***), the first principal axis of the protein complex is parallel to the *z*-axis. We then embedded the aligned TCR-CD3 complex with the CHARMM-GUI program (***Wu et al., 2014***; ***Jo et al., 2008***; ***Lee et al., 2020***) into a lipid membrane that is oriented along the *x*-*y*-plane of the simulation box and is composed of 228 palmitoyl-2-oleoyl-sn-glycero-3-phosphocholine (POPC) and 57 cholesterol molecules in each monolayer, added missing atoms of the proteins with this program, and capped the N− and C-terminal ends of the eight protein chains with neutral ACE (−COCH_3_) and NME (−NHCH_3_) residues. We solvated this membrane-protein assembly at a salt concentration of 0.15 M KCl such that a 2.5 nm thick water layer is maintained above and below in *z*-direction. We performed the membrane embedding and solvation ten times to obtain ten system conformations as starting conformations of our simulations. The number of water molecules in this ten conformations slightly varies from 79 744 to 79 855.

### System equilibration

We have equilibrated the ten system conformations with the Amber16 software (***Case et al., 2017***). In this equilibration, we have first performed an energy minimization with 5000 minimization steps of steepest decent and subsequent 5000 steps of the conjugent gradient algorithm. The positions of backbone atoms of the EC and TM domains of the proteins were harmonically restrained in this minimization with a force constant of 10 kcal mol^−1^ Å^−2^. We have subsequently heated the systems in two simulation steps of 5 ps and 100 ps with harmonic restraints on all protein and lipid atoms: (1) from 0 K to 100 K at constant volume, and (2) from 100 K to 303 K at a constant pressure of 1 bar using the Berendsen barostat with semi-isotopic pressure coupling and a pressure relaxation time of 2 ps. In both heating steps, we used a Langevin thermostat with a collision frequency of 1 ps−1, a MD integration time step of 2 fs, and a force constant of 10 kcal mol^−1^ Å^−2^ for the harmonic restraints. We have finally performed equilibration simulations with a total length of 20 ns at the temperature 303 K and a constant pressure of 1 bar. The equilibration simulations were carried out in ten steps of 2 ns with decreasing harmonic restraints on the protein backbone atoms of 10.0, 8.0, 6.0, 4.0, 2.0, 0.8, 0.6, 0.4, 0.2, and 0.1 kcal mol^−1^ Å^−2^ in these steps. We used a Langevin thermostat with collision frequency 1.0 ps^−1^, a Berendsen barostat with a pressure relaxation time of 1 ps for the semi-isotropic pressure coupling, and in integration time step of 2 fs in these simulations. The lengths of all bond involving hydrogens were constrained with the SHAKE algorithm (***Miyamoto and Kollman, 1992***; ***Ryckaert et al., 1977***), and a cutoff length of 1.0 nm was used in calculating the non-bonded interactions with the Particle Mesh Ewald (PME) algorithm ***Essmann et al. (1995***); ***Darden et al. (1993***).

### Production simulations

For each of the ten equilibrated system conformations, we have generated 12 independent MD trajectories with a length of 1 μs at a temperature of 303 K. The total simulation time of these 120 production trajectories thus is 120 μs. We conducted the production simulations with the Amber99SB-ILDN protein force field (***Lindorff-Larsen et al., 2010***), the Amber Lipid14 membrane force field (***Dickson et al., 2014***), and the software AMBER 16 GPU (***Salomon-Ferrer et al., 2013***; ***Le Grand et al., 2013***). We used a Langevin thermostat with a Langevin collision frequency of 1.0 ps−1, and the Berendsen barostat with semi-isotropic pressure coupling and a relaxation time of 1 ps to apply a constant pressure of 1 bar in all directions at which the membrane is tensionless. As in the equilibration simulations, the lengths of all bond involving hydrogens were constrained with the SHAKE algorithm (***Miyamoto and Kollman, 1992***; ***Ryckaert et al., 1977***), and a cutoff length of 1.0 nm was used in calculating the non-bonded interactions with the Particle Mesh Ewald (PME) algorithm (***Essmann et al., 1995***; ***Darden et al., 1993***). In addition, we applied hydrogen mass repartitioning (***Hopkins et al., 2015***) to all the hydrogen atoms on protein and lipids in the production simulations, which allowed to increase to the MD integration time step to 4 fs.

### Analysis of trajectories

Our analysis of the structural and orientational variability of the TCR - CD3 complex along the simulation trajectories is based on the residue-residue contacts of the TCRαβ, CD3єδ, and CD3єγ EC domains, on the tilt and rotation angle of the TCRαβ EC domain relative the membrane plane, and on the inclination angles of the TM helices relative the membrane. We take two residues from different EC domains to be in contact if the minimum distance between non-hydrogen atoms of the residues is smaller than 0.45 nm. Our tilt and rotation angles of the TCRαβ EC domain are determined from two characteristic axes in the domain: Axis A connects the centres of mass of Cαβ and Vαβ, where Cαβ is the dimer of the constant domains Cα and Cβ, and Vαβ is the dimer of the variable domains Vα and Vβ. Axis B connects the centres of mass of the variable domains Vα and Vβ. We define the tilt angle of the TCRαβ EC domain as the angle between axis A and the membrane normal, and the rotation angle as the angle between axis B and the normal of the plane spanned by axis A and the membrane normal. The rotation angle describes the rotation of the TCRαβ EC domain around axis A. To determine the inclination angles of the TM helices, we divide the residue span of the helices defined in the PDB file 6jxr of the cryo-EM structure (***Dong et al., 2019***) into two halves. We calculate the inclination angle of a TM helix as the angle between the membrane normal and the line that connects the centres of mass of the two helix halves.

Figure 4 indicates that the structural and orientational ensembles sampled by our 120 production trajectories equilibrate within the first 0.5 μs of the trajectories. The data points in the figure represent averages over the 120 simulation structures of the trajectories at the indicated simulation times, with error bars representing the error of the mean for these 120 frames. The time-dependent trajectory averages for the tilt and rotation angle Figure 4(a) converge to average values of about 26° for the rotation angle and between 33° and 34° for the tilt angle within 0.5 μs. Within this time, the inclination angles of TM helices in Figure 4(b) and the numbers of contacts for structural elements in the TCR constant domains Cα and Cβ and for the variable domain Vα converge as well. We focus in our analysis therefore on the second trajectory halves and extract 50 structures at the simulations times 0.51 μs, 0.52 μs, 0.53 μs,…, 1.0 μs from each trajectory, which results in 120 × 50 = 6000 structures that reflect an equilibrated ensemble of simulation conformations. The contacts, contact numbers, and angles presented in Figures 2 and 3 are calculated from these 6000 structures. The 6000 structures are available at the Edmond Open Research Data Repository at https://dx.doi.org/10.17617/3.5m (***Pandey and Weikl, 2021***).

**Figure 4.**
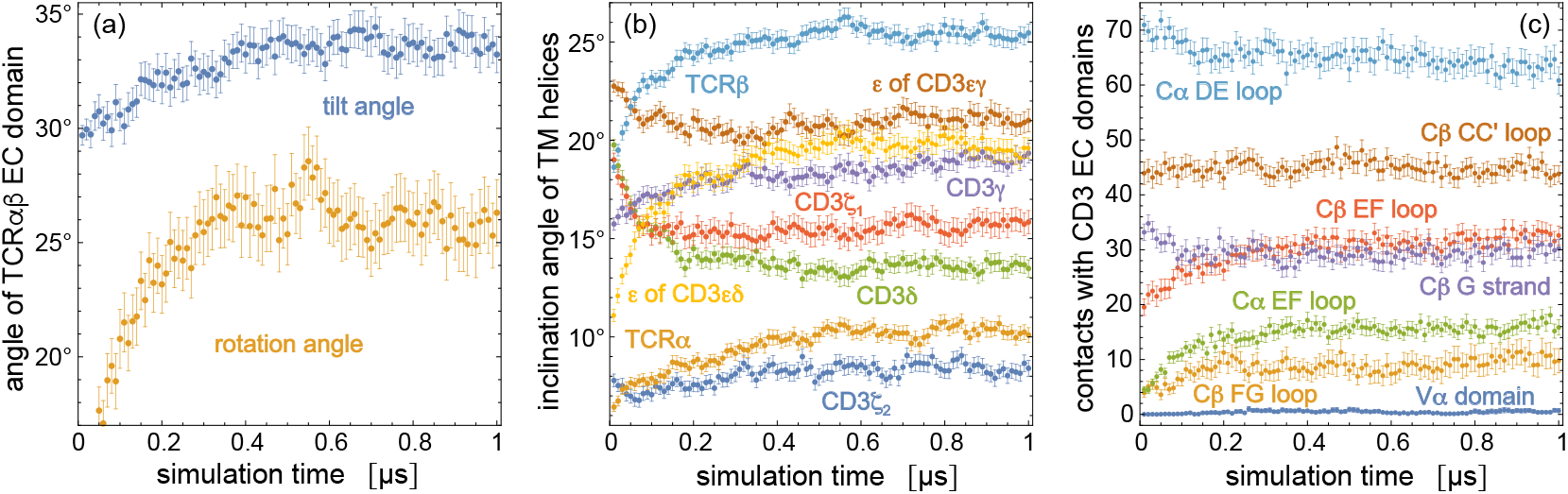
Time-dependent trajectory averages for (a) the tilt angle and rotation angle of the TCRαβ EC domain, (b) the inclination angles of the TM helices of the eight protein chains relative to the membrane normal, (c) the number of contacts of structural elements in the TCR constant domains Cα and Cβ and of the variable domain Vα with the two CD3 EC domains. Each data point is an average over the simulation structures of the 120 trajectories at the indicated time point, with error bars representing the error of the mean for these 120 structures. The structural elements of Cα and Cβ are de1ned in Figure 2.

---

**Figure 2–Figure supplement 1.**
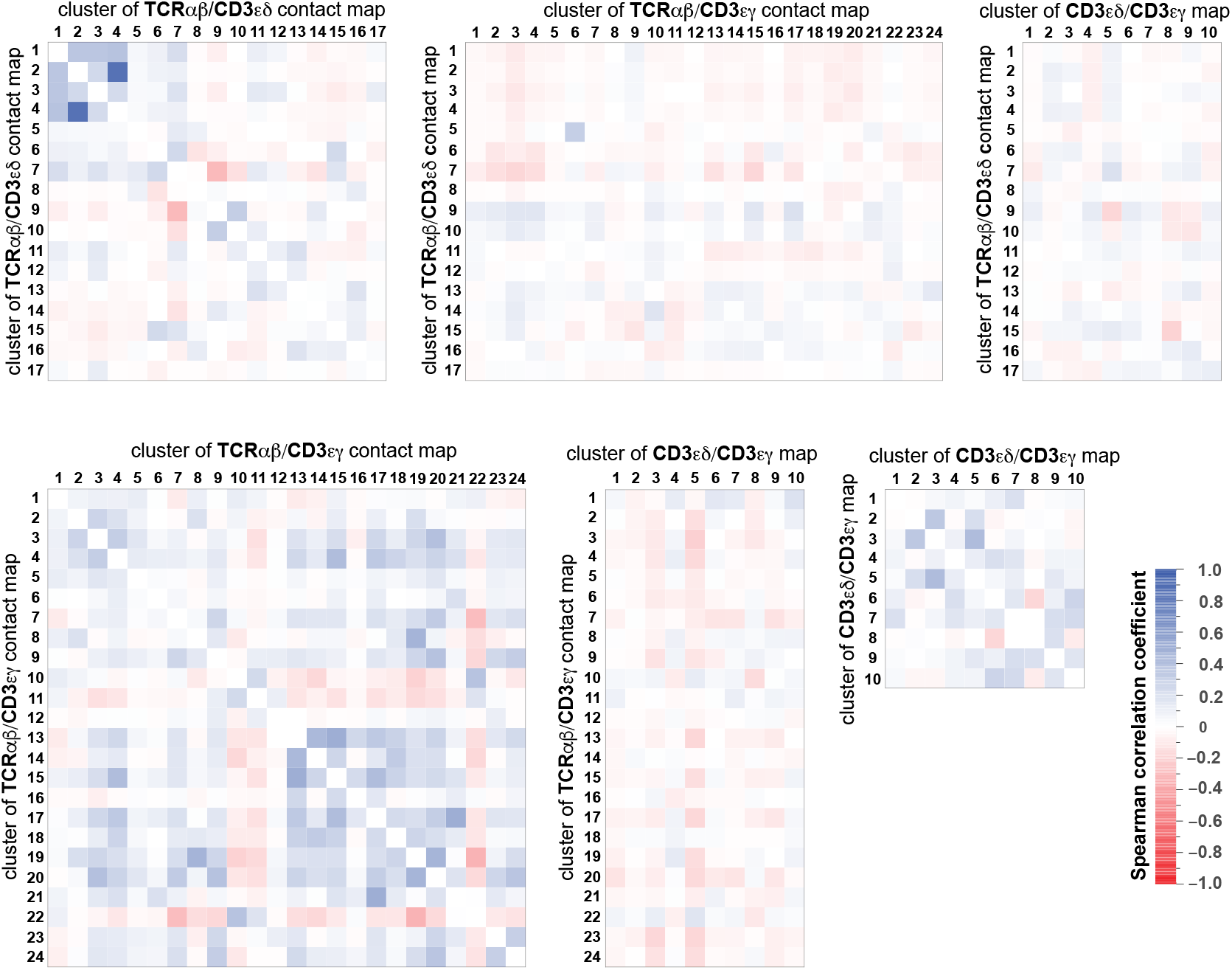
Spearman correlation cofficients of the contact clusters in the contact maps of Figure 2. The Spearman correlation coefficient assesses monotonic relations that can be linear or nonlinear, and is calculated here for the numbers of contacts between non-hydrogen atoms of each cluster in the 6000 simulation frames of the contact analysis. the MD simulation trajectories in our analysis.

**Figure 3–Figure supplement 1.**
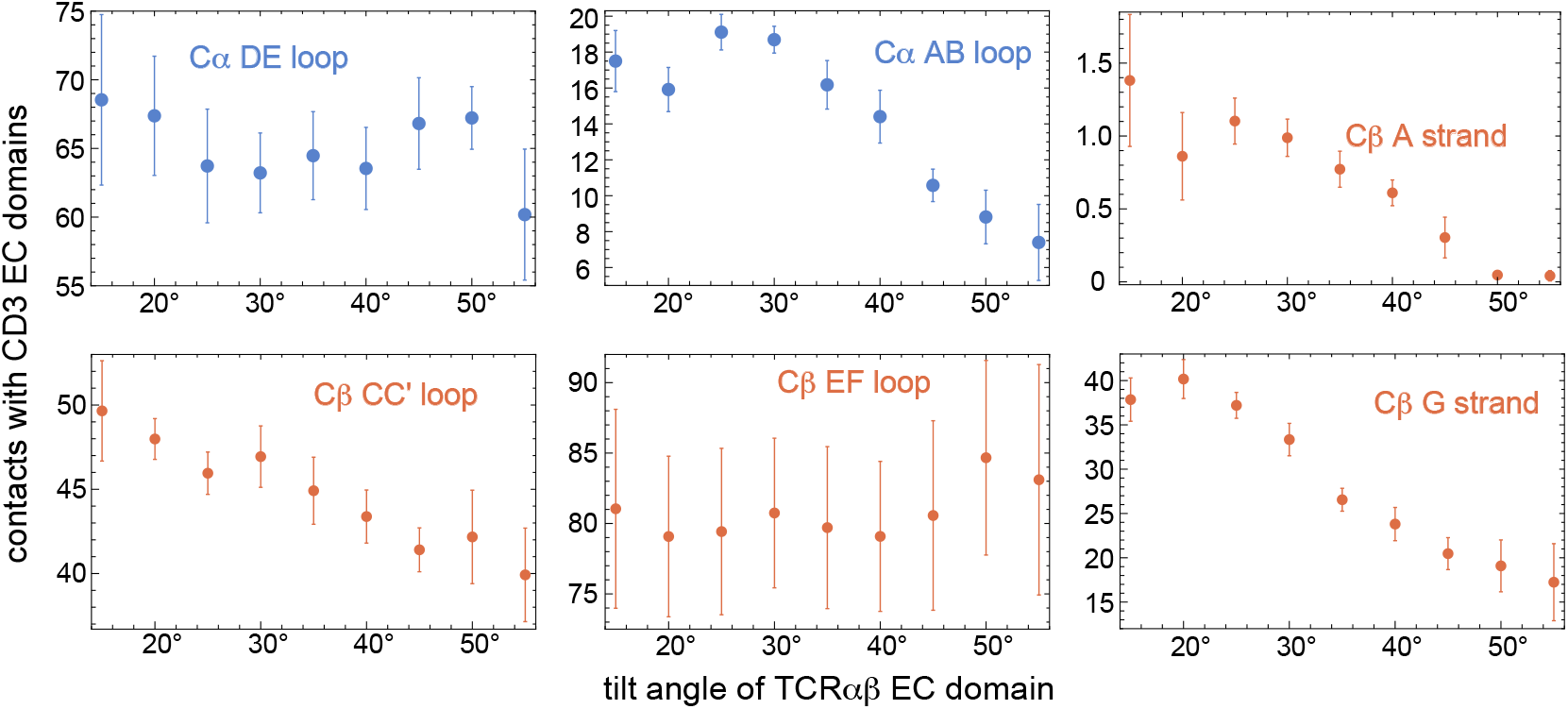
Numbers of residue-residue contacts of TCR Cα and Cβ loops and strands in interaction with CD3s versus tilt angle of the TCRαβ EC domain.

**Figure 3–Figure supplement 2.**
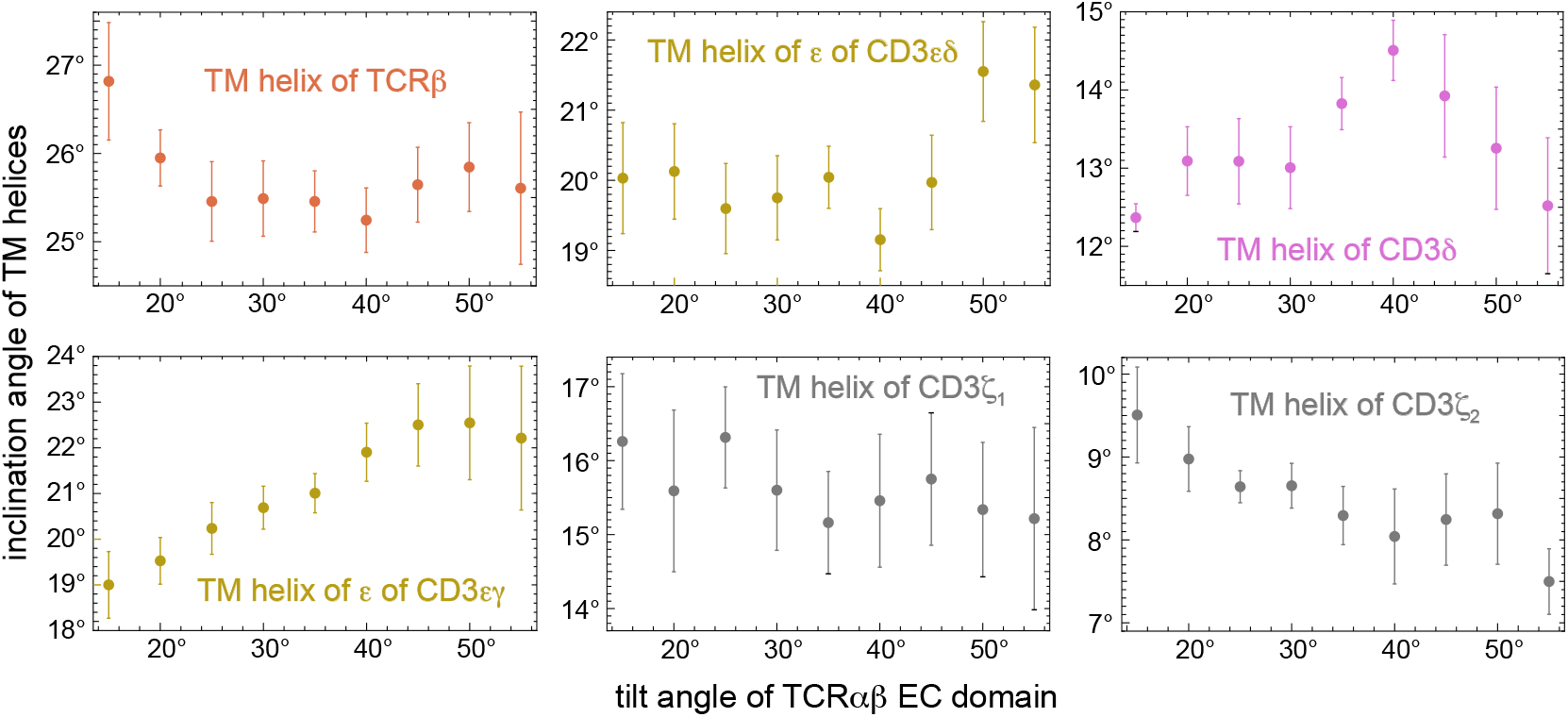
Inclination angles of TM helices relative to the membrane normal as a function of the tilt angle of the TCRαβ ECdomain.

